# Ratiometric growth-rate control enables robust coexistence in competing microbial consortia

**DOI:** 10.64898/2026.08.28.747825

**Authors:** Carlos Barajas

## Abstract

Maintaining a prescribed composition in engineered microbial consortia is difficult because small fitness differences can drive competitive exclusion. We study a two-strain consortium in continuous culture and develop a feedback architecture that regulates composition by selectively slowing the fast strain as a function of the population ratio. At the population level, we derive an idealized ratio-feedback law with a tunable positive coexistence equilibrium. We then propose a biomolecular realization using orthogonal quorum sensing, an sRNA-based ratiometric controller, and a ppGpp-mediated growth actuator. Exploiting the separation between slow population growth and faster intracellular controller dynamics, we use singular perturbation theory to show that, for sufficiently fast controller dynamics, the full implementation model inherits the coexistence equilibrium and its local stability properties from the reduced model. Numerical simulations validate the reduction and show how weaker timescale separation or loss of the assumed molecular regime degrades performance.

## I. Introduction

Engineered microbial consortia are attractive for bioproduction, biosensing, therapeutics, and environmental remediation because they enable division of labor and reduce the burden on individual strains [1]–[4]. In practice, however, these benefits depend on maintaining a desired population composition. When strain growth rates are mismatched, even small fitness differences are amplified over time, leading to competitive exclusion, especially in continuous culture [5], [6].

A range of strategies has been proposed to regulate microbial communities, including cybergenetic control, reactor-level architectures, and genetically embedded feedback acting through quorum sensing and growth modulation [7]–[11]. Beyond coexistence, many applications require regulation to a prescribed population ratio, for example to enforce division of labor or pathway stoichiometry [1], [4].

Ratiometric sensing and control principles have appeared in natural and engineered settings, including population sensing, phenotype regulation, intracellular coculture control, and dual-chamber bioreactor control [8], [12]–[14]. However, for intracellular composition controllers, what is still missing is a control-theoretic framework to analyze the design, identify which parameters are tunable for programmable and stable operation, and characterize the operating regimes in which the design works as intended.

This paper addresses that gap for a continuously operated two-strain consortium in which one strain grows faster than the other. We first derive an idealized ratio-feedback law that guarantees a tunable positive coexistence equilibrium at the population level. We then propose a modular biomolecular realization (Fig. 1) based on orthogonal quorum sensing [15], an sRNA-based ratiometric sensor [14], [16], and a ppGpp-mediated growth actuator [17]. Exploiting the separation between slow population growth and faster intracellular controller dynamics, we use singular perturbation theory to show that, under explicit operating-regime assumptions, the full implementation model inherits the coexistence equilibrium and its local stability properties from the reduced model. Numerical simulations validate the reduction and show how violations of these assumptions degrade performance.

**Fig. 1.**
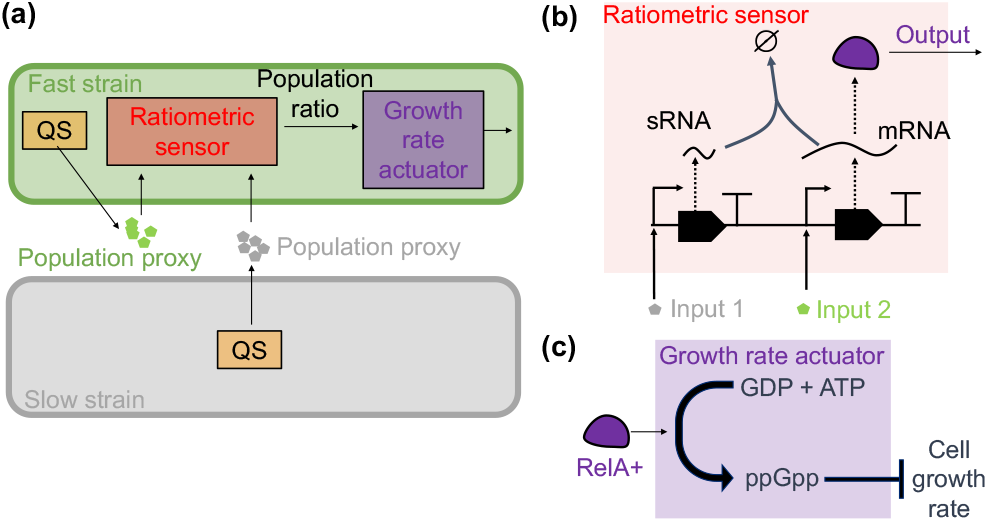
Proposed modular implementation for ratio regulation in a two-strain consortium. (a) Closed-loop architecture: quorum-sensing (QS) channels provide population proxies to a ratiometric controller in the fast strain, whose output drives a growth-rate actuator to modulate fast-strain fitness. (b) Ratiometric controller module: the slow-strain QS signal (Input 1) induces expression of an sRNA that degrades an mRNA whose expression is driven by the fast-strain QS signal (Input 2). The mRNA encodes the output protein, whose concentration is proportional to the ratio of Input 2 to Input 1. (c) Growth-rate actuator module: RelA-mediated ppGpp production represses cell growth, providing the actuation mechanism for regulating population composition.

## II. Population Dynamics and Control Objective

We consider a well-mixed chemostat containing two strains. The slow strain, with concentration *X*_*s*_, and the fast strain, with concentration *X*_*f*_, compete for a shared carrying capacity *K >* 0 and are diluted at rate *D >* 0. The population dynamics are

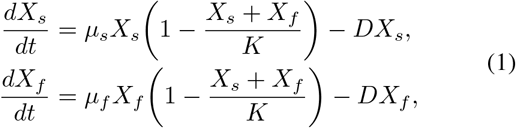

where *µ*_*s*_ *>* 0 and *µ*_*f*_ *>* 0 are the intrinsic growth rates of the slow and fast strains, respectively. In open loop we assume *µ*_*f*_ *> µ*_*s*_, so that the fast strain has a competitive advantage in the absence of feedback.

Define the normalized states and time

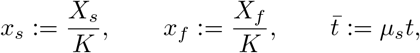

together with the dimensionless parameters

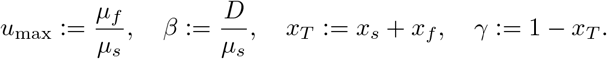

The normalized dynamics become

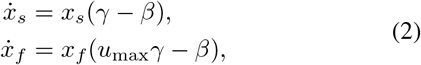

where henceforth 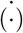 denotes differentiation with respect to normalized time 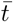. The quantity *x*_*T*_ = *x*_*s*_ + *x*_*f*_ is the total normalized population, and *γ* = 1 − *x*_*T*_ is the remaining fraction of unused carrying capacity. The total population satisfies

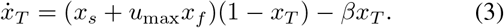

### A. Open-loop competitive exclusion

For the ratio

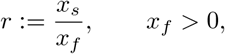

(2) gives

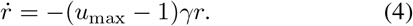

Thus the ratio decays whenever *u*_max_ *>* 1. Moreover, if *x*_*T*_ (0) *<* 1 and 0 *< β < u*_max_, then

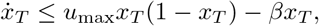

so comparison gives

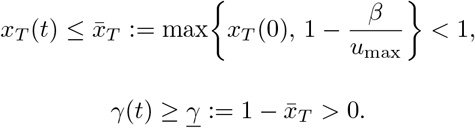

Hence

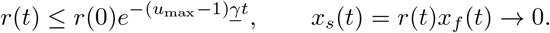

Open-loop coexistence is therefore lost: the fast strain excludes the slow strain. The control objective is to restore coexistence by regulating the ratio *r* = *x*_*s*_*/x*_*f*_ to a tunable positive setpoint *r*_ss_ *>* 0 through feedback on the growth rate of the fast strain.

## III. CLOSED-LOOP RATIO REGULATION FRAMEWORK

To regulate composition, replace the constant growth-rate ratio *u*_max_ by an idealized ratio-feedback law

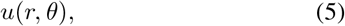

where *θ* sets the desired steady-state ratio.

### Assumption 1.

*For each fixed θ, the map u*(·, *θ*) : (0, ∞) → (0, ∞) *is C*^1^, *strictly increasing, and bounded:*

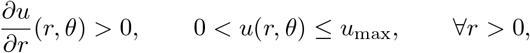

*for some constant u*_max_ *>* 1. *In addition, there exist r*_−_, *r*_+_ *>* 0 *such that*

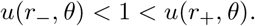

Replacing *u*_max_ in (4) by *u*(*r, θ*) gives

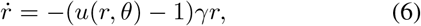

so any positive equilibrium ratio must satisfy

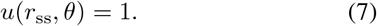

By Assumption 1, this equation has a unique solution *r*_ss_ *>* 0.

### Theorem 1.

*Suppose Assumption 1 holds, x*_*T*_ (0) *<* 1, *r*(0) *>* 0, *and* 0 *< β < u*_max_. *Then* (7) *admits a unique solution *r*_*ss*_ > 0. Define*

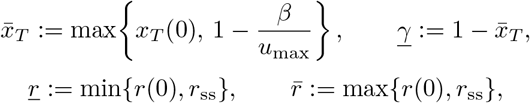

*and*

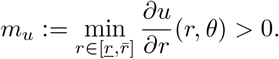

*Then* 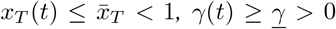, *the interval* 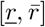 *is forward invariant, and*

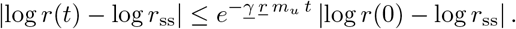

*In particular, r*(*t*) → *r*_ss_.

*Proof*. Existence and uniqueness of *r*_ss_ follow from continuity, strict monotonicity, and the crossing condition in Assumption 1. For the total population,

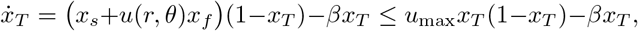

so the same comparison argument as in the open-loop case gives the bounds on *x*_*T*_ and *γ*. Since *u*(·, *θ*) is strictly increasing and *u*(*r*_ss_, *θ*) = 1,

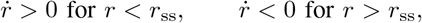

which gives forward invariance of 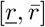. Let *ρ* := log *r* and *ρ*_ss_ := log *r*_ss_. Then

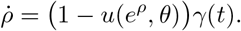

By the mean value theorem,

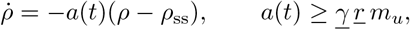

on the invariant interval 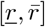. Gronwall’s inequality yields the stated estimate.

Thus the design task is to realize an idealized ratio-feedback law *u*(*r, θ*) that is increasing in *r*, bounded, and crosses unity at the desired ratio. The parameter *θ* then directly determines the steady-state composition.

## IV. Biological Realization of Ratiometric Growth-Rate Feedback

We now derive a biomolecular realization of the idealized ratio-feedback law *u*(*r, θ*) introduced above. The implementation is organized into three modules: a quorum-sensing layer that encodes strain abundance, an sRNA-based ratiometric controller that computes a ratio-dependent intracellular signal, and a ppGpp-based actuator that modulates the growth rate of the fast strain.

### A. Quorum-sensing for population size proxy

Each strain produces an orthogonal quorum-sensing molecule that serves as a proxy for its population size (Fig. 1a). Let *q*_*s*_ and *q*_*f*_ denote the effective intracellular concentrations of the signals associated with the slow and fast strains, respectively. We assume that diffusion and transport between the extracellular medium and the cell interior are fast relative to cell growth, so that intracellular and extracellular signal levels are well-mixed and can be represented by a single effective concentration.

A minimal model is

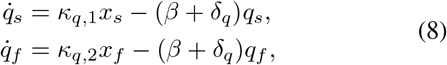

where *κ*_*q*,1_ and *κ*_*q*,2_ lump together production, diffusion, and uptake, and *δ*_*q*_ represents signal removal through degradation or quorum quenching (e.g., enzymatic degradation of Acyl-homoserine lactone signals by lactonases).

### B. sRNA-mediated ratiometric controller

We next construct an intracellular module that transforms the two quorum-sensing inputs into a ratio-dependent actuator level. Let *m*_*A*_ denote the mRNA encoding an actuator protein *A*, and let *S* denote an sRNA that promotes degradation of *m*_*A*_. The controller dynamics are

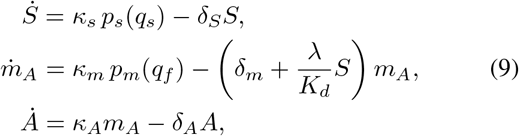

where *κ*_*s*_ and *κ*_*m*_ are maximal transcription rates, *κ*_*A*_ is the translation rate of *A*, and *δ*_*S*_, *δ*_*m*_, and *δ*_*A*_ denote degradation (and dilution) rates of *S, m*_*A*_, and *A*, respectively. The term (*λ/K*_*d*_)*S* captures an effective sRNA-mediated degradation rate of *m*_*A*_, where *λ* is an interaction rate constant and *K*_*d*_ is an effective binding or sequestration constant. For simplicity, we assume that the sRNA pool is maintained in excess relative to *m*_A_, so that depletion of *S* through binding does not explicitly appear in the *S* dynamics. The functions *p*_*s*_(·) and *p*_*m*_(·) represent regulated transcription driven by the quorum-sensing signals *q*_*s*_ and *q*_*f*_, respectively, which we approximate using Hill-type functions

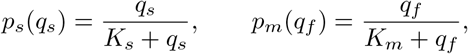

where *K*_*s*_ and *K*_*m*_ are effective dissociation constants.

### C. ppGpp-mediated growth-rate actuation

The controller output *A* is used to modulate growth of the fast strain through the ppGpp pathway. Specifically, we employ a RelA^+^ actuator, a truncated version of RelA that retains ppGpp synthesis activity but lacks regulatory domains, and has been previously used to dynamically control cell growth in engineered systems [17]. Expression of RelA^+^ increases intracellular ppGpp levels, which in turn repress cellular growth.

We describe this effect by

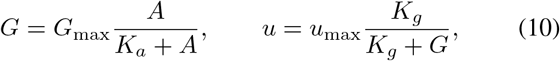

where *G* denotes the effective ppGpp-mediated growth inhibition level, *G*_max_ is the maximal ppGpp load, *K*_*a*_ is the activation constant relating actuator expression to ppGpp production, and *K*_*g*_ sets the sensitivity of growth rate to ppGpp.

### D. Quasi-steady-state reduction to the idealized ratio-feedback law

We now show how the three biological modules combine to approximate the idealized ratio-feedback law *u*(*r, θ*). Assume that the molecular states *q*_*s*_, *q*_*f*_, *S, m*_*A*_, and *A* evolve on a faster timescale than the population states *x*_*s*_ and *x*_*f*_. Setting the molecular dynamics to quasi-steady state with *x*_*s*_ and *x*_*f*_ held fixed gives

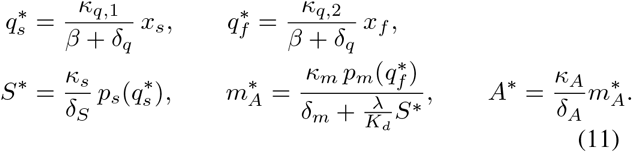

To obtain a transparent expression, consider the operating regime in which

i. the promoters operate in their linear range,

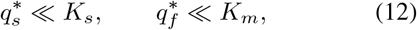

so that

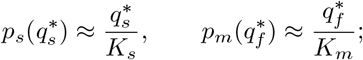
ii. sRNA-mediated removal dominates basal mRNA decay,

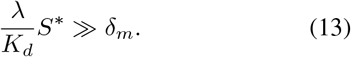

Under these assumptions,

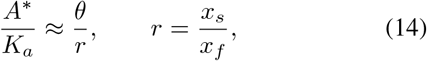

where

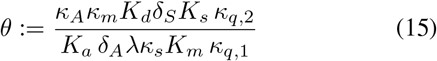

is a lumped design parameter determined by the circuit parameters.

Substituting (14) into (10) yields

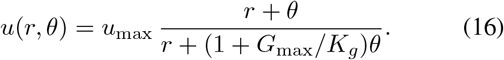

For *θ >* 0 and *G*_max_*/K*_*g*_ *>* 0, this law is *C*^1^, positive, bounded, and strictly increasing in *r*. Moreover,

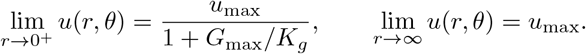

Hence, if

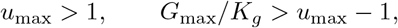

then *u*(*r, θ*) crosses unity exactly once and satisfies Assumption 1. Biologically, this requires that ppGpp-mediated repression be strong enough to drive the effective growth rate of the fast strain below that of the slow strain when needed. Unlike *θ*, which serves as a tuning parameter for the target ratio, this is a feasibility condition determined by the achievable host-circuit operating regime. This should be interpreted as a nominal feasibility condition for the idealized ratio-feedback law. In the full implementation model, however, the achievable actuator level is finite because *x*_*T*_ ≤1− *β* and the molecular modules saturate.

The steady-state ratio is determined by the unity-crossing condition *u*(*r*_ss_, *θ*) = 1, which yields

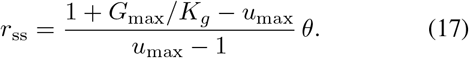

Thus, *θ* acts as a direct tuning parameter for the equilibrium population ratio. In practice, *θ* can be adjusted through circuit design parameters, including the relative strengths of the quorum-sensing promoters, the sRNA–mRNA interaction rate, and the expression level of the RelA^+^ actuator.

This tuning rule is sensitive to the feasibility margin

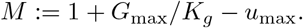

Since *r*_ss_ = *θM/*(*u*_max_ − 1), uncertainty in either the achievable ppGpp inhibition *G*_max_*/K*_*g*_ or the open-loop growth advantage *u*_max_ is amplified when *M* is small. Thus practical designs should not operate close to the boundary *G*_max_*/K*_*g*_ = *u*_max_ −1. Instead, the actuator should provide sufficient margin to tolerate host-dependent variation in growth burden, ppGpp sensitivity, and basal growth rate.

## V. Singular Perturbation Analysis of the Full Controller

The previous section derived the idealized ratio-feedback law *u*(*r, θ*) from the biomolecular implementation under a quasi-steady-state approximation. We now analyze the full implementation model and justify this reduction using singular perturbation theory. We proceed in three steps: we first establish boundary-layer stability, then analyze the reduced coexistence equilibrium, and finally transfer these properties to the full implementation model for sufficiently small *ϵ*.

We write the full implementation model in standard singular perturbation form

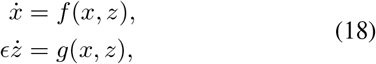

Where

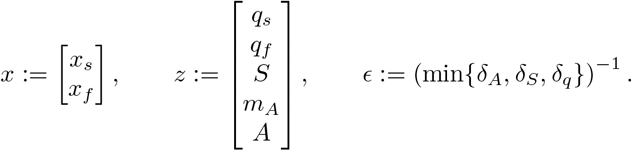

The slow vector field is

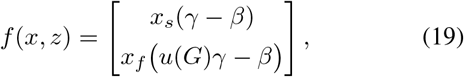

and the fast vector field is

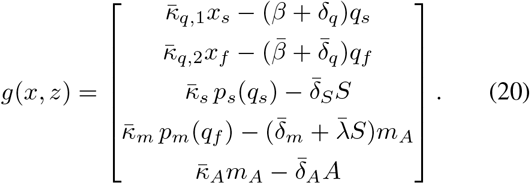

Here,

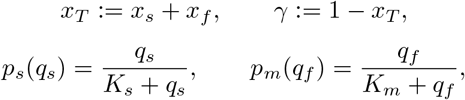

and the ppGpp-mediated growth feedback is

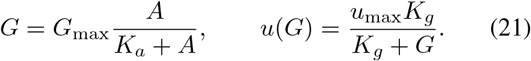

An overbar denotes the corresponding parameter normalized with respect to the fast timescale.

## A. Boundary-layer dynamics and stability

Treating *x* as frozen, the algebraic equation *g*(*x, z*) = 0 defines the quasi-steady-state map

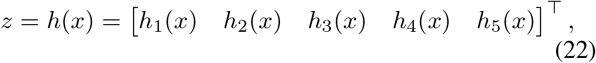

where

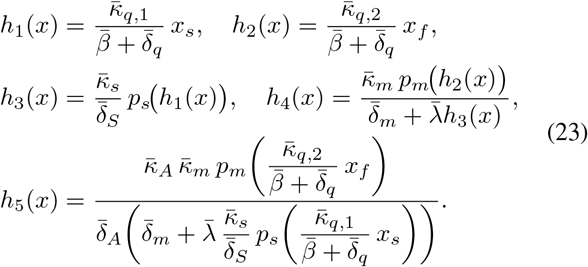

Define the boundary-layer coordinate

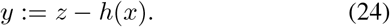

With stretched time *τ* = *t/ϵ*, the fast dynamics become

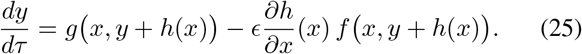

Setting *ϵ* = 0 with *x* frozen gives the boundary-layer model

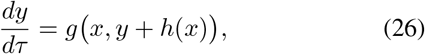

whose equilibrium is *y* = 0.

Linearizing (26) about *y* = 0 yields a lower-triangular Jacobian with diagonal entries

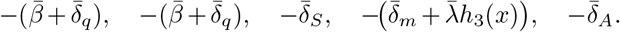

Since all parameters are positive and *h*_3_(*x*) ≥ 0, these entries are uniformly bounded away from the imaginary axis. Hence, for each fixed *x*, the equilibrium *y* = 0 of (26) is locally exponentially stable, uniformly on compact subsets of the slow-state space.

### B. Reduced system, coexistence equilibrium, and stability

Substituting *z* = *h*(*x*) into the slow dynamics gives the reduced system

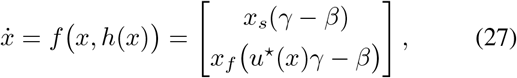

where

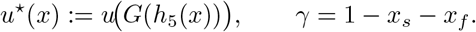

Let

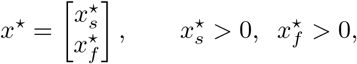

denote a coexistence equilibrium of (27). Since both components are positive, the equilibrium conditions are

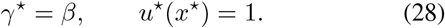

The first relation gives

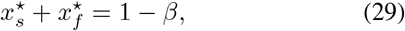

so a necessary condition is

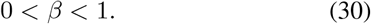

Define

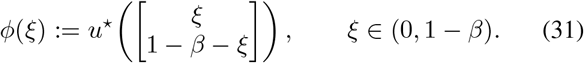

Since *h*_5_ decreases with *x*_*s*_ and increases with *x*_*f*_, while *G*(·) is increasing and *u*(·) is decreasing, we have

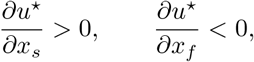

and hence

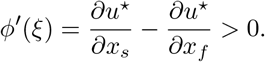

Thus *ϕ* is strictly increasing, and a positive coexistence equilibrium exists if and only if

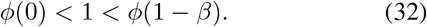

At *ξ* = 1−*β*, we have *x*_*f*_ = 0, so *h*_2_ = *h*_4_ = *h*_5_ = 0, and therefore

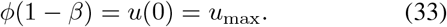

At *ξ* = 0, let

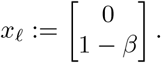

Then

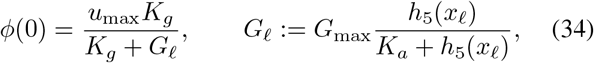

with

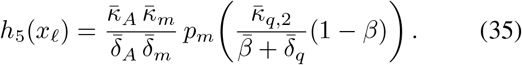

Hence (32) is equivalent to

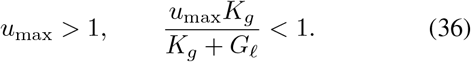

Under (30) and (36), the reduced system admits a unique positive coexistence equilibrium. Biologically, (36) requires that ppGpp-mediated repression be strong enough to drive the effective growth rate of the fast strain below that of the slow strain under maximal activation.

The Jacobian of (27) at *x*^*^ is

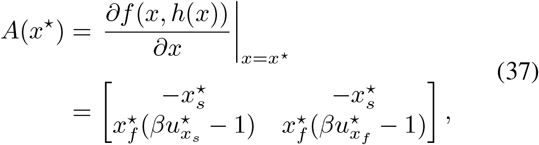

where

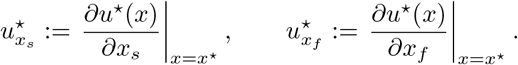

Since 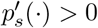 and 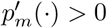, the map *h*_5_(*x*) satisfies

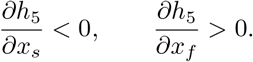

Together with 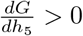 and 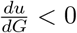, the chain rule gives

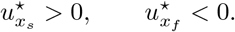

Hence

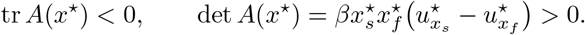

Therefore *A*(*x*^*^) is Hurwitz, and the coexistence equilibrium *x*^*^ is locally exponentially stable.

### C. Approximation of the idealized ratio-feedback law

#### Theorem 2.

*Let* (*x*(*t, ϵ*), *z*(*t, ϵ*)) *denote the solution of* (18) *with*

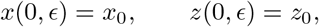

*and let x*(*t*) *denote the solution of the reduced system* (27) *with x*(0) = *x*_0_. *Let y*(*τ*) *denote the solution of the frozen boundary-layer problem*

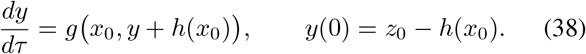

*Assume x*_0_ *belongs to a compact subset of the region of attraction of the reduced equilibrium x*^*^, *and z*_0_ − *h*(*x*_0_) *belongs to a compact subset of the region of attraction of y* = 0 *for* (38). *Then there exists ϵ*^*^ *>* 0 *such that, for all* 0 *< ϵ < ϵ*^*^,

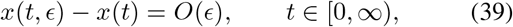

*and*

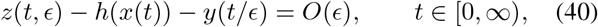

*uniformly on the infinite interval. Moreover, for any t*_b_ *>* 0,

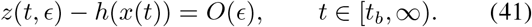

*Proof*. After translating *x*^*^ to the origin, the reduced system has an exponentially stable equilibrium and the boundary-layer system has an exponentially stable equilibrium *y* = 0, uniformly in *x*, by the preceding analysis. Since *f, g*, and *h* are smooth, the hypotheses of [18, Thm. 11.2] are satisfied locally. The conclusions then follow directly.

#### Remark 1.

*In addition to the conditions of Theorem 2, if the operating-regime assumptions* (12)*–*(13) *hold near the reduced equilibrium x*^*^, *then the effective growth law of the full implementation model satisfies*

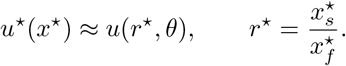

*Thus, near equilibrium, the full implementation model realizes the idealized ratio-feedback law*.

### D. Full closed-loop stability for small ϵ

We now transfer the reduced and boundary-layer stability properties to the full implementation model.

#### Theorem 3.

*Assume* (30) *and* (36) *hold. Then the full implementation model* (18) *has the equilibrium*

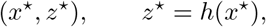

*where x*^*^ *is the unique positive equilibrium of the reduced system* (27). *Moreover, there exists* 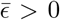 *such that, for all* 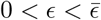 *this equilibrium is locally exponentially stable*.

*Proof*. By the preceding reduced-order analysis, (30) and (36) guarantee that (27) admits a unique positive equilibrium *x*^*^. Since *z*^*^ = *h*(*x*^*^) satisfies

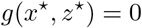

by definition of the quasi-steady-state map, and

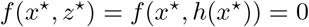

because *x*^*^ is an equilibrium of the reduced system, it follows that (*x*^*^, *z*^*^) is an equilibrium of the full implementation model.

Translate (*x*^*^, *z*^*^) to the origin and introduce the boundary-layer coordinate *y* = *z*− *h*(*x*). By the preceding subsections, the shifted reduced system has an exponentially stable equilibrium at the origin, and the boundary-layer system has an exponentially stable equilibrium *y* = 0, uniformly in the frozen slow variable. Since *f, g*, and *h* are smooth, the local hypotheses of [18, Thm. 11.4] are satisfied after translation. Hence, for sufficiently small *ϵ*, the origin of the shifted full implementation model is exponentially stable, which is equivalent to local exponential stability of (*x*^*^, *h*(*x*^*^)) in the original coordinates.

#### Remark 2.

*Theorem 3 shows that the feedback controller induces coexistence: the full implementation model admits a locally exponentially stable equilibrium with both strains present, thereby enforcing the desired population composition despite intrinsic competitive exclusion*.

Theorem 1 gives global ratio convergence for the idealized two-state law, whereas Theorem 3 gives only local exponential stability for the full biomolecular implementation. This locality is inherited from the singular-perturbation argument, which applies on compact subsets of the attraction regions of the reduced and boundary-layer systems. Consequently, the full-system basin of attraction may depend on *ϵ* and the initial distance from *z* = *h*(*x*).

## VI. Numerical Simulations

The numerical results make two points. First, the full implementation model (18) can reproduce the idealized ratio-feedback law obtained by replacing *u*_max_ in (2) with (16). Second, this agreement requires both small *ϵ* and the operating-regime assumptions (12)–(13) used to derive the ratiometric relation.

Figure 2 compares the idealized two-state model (2) with *u*_max_ replaced by (16) against the full implementation model (18). When *ϵ* is small and (12)–(13) hold, the full implementation model reproduces the same setpoint regulation as the idealized two-state model. This is observed for *ϵ* = 0.1, where trajectories match the predicted monotone convergence. For larger *ϵ*, timescale separation is reduced and the feedback behaves with delay, leading to overshoot and oscillatory transients while still regulating the composition.

**Fig. 2.**
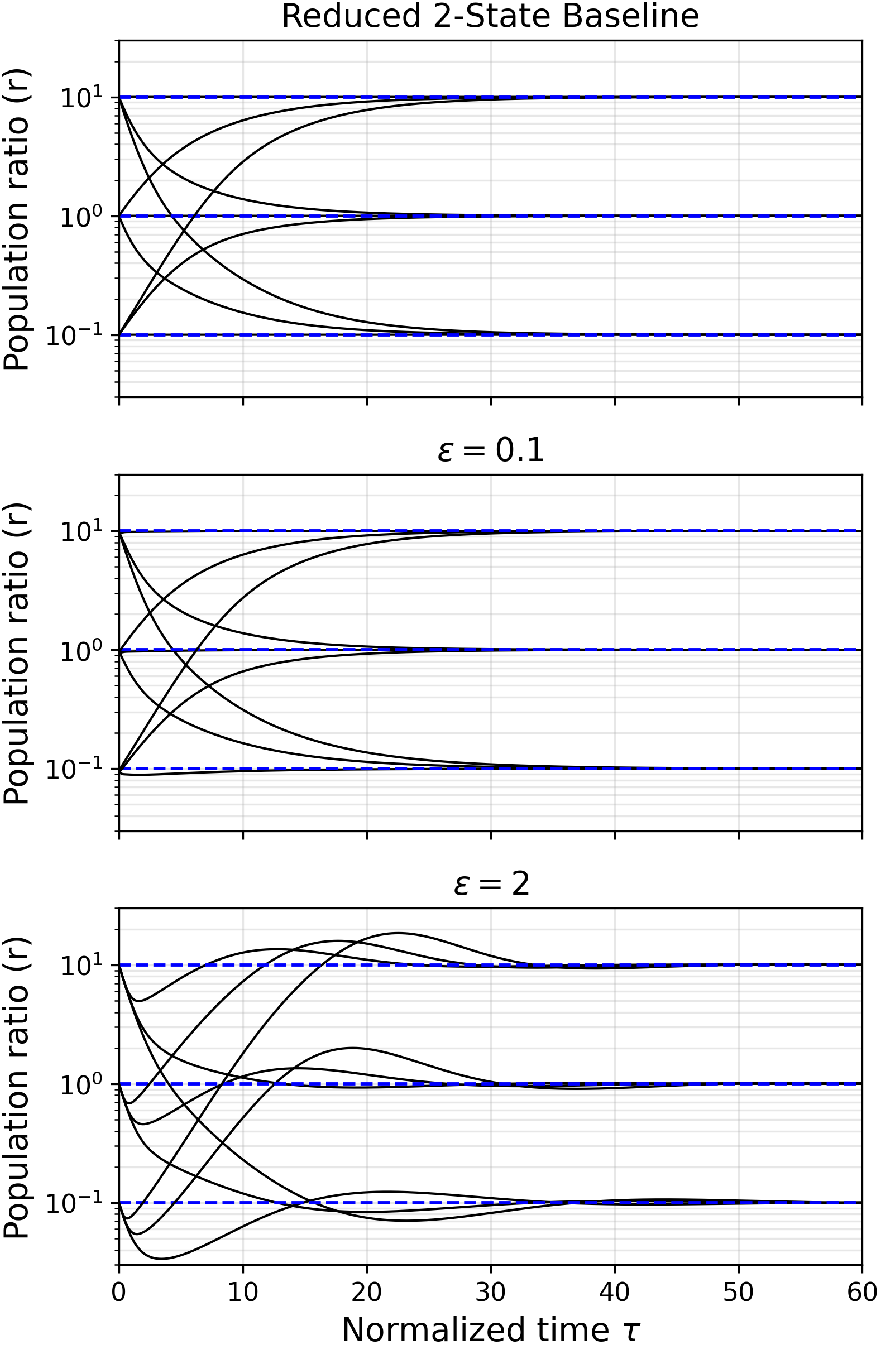
When the implementation assumptions hold, the full implementation model reproduces the idealized ratio-feedback law; increasing. *ϵ* **introduces delay-like oscillatory transients**. The top panel simulates the idealized two-state model (2) with *u*_max_ replaced by (16); the lower panels simulate the full implementation model (18). Parameters: 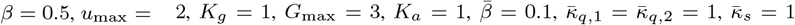, 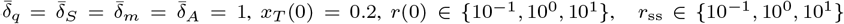, and tuned 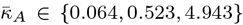 Initial states in all full-model runs: *x*_*s*_(0) = *x*_*T*_ (0)*r*(0)*/*(1 + *r*(0)), *x*_*f*_ (0) = *x*_*T*_ (0)*/*(1 + *r*(0)), *q*_*s*_(0) = *q*_*f*_ (0) = *S*(0) = *m*_*A*_(0) = *A*(0) = 0. The parameter choices satisfy the assumptions used to derive (16): conditions (12) and (13) hold, 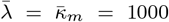 makes sRNA-mediated mRNA removal dominant, and *K*_*s*_= *K*_*m*_ = 10 keeps the promoter maps close to their linear regime over the simulated trajectories. The top panel uses *θ* ∈ {0.05, 0.5, 5 }. The middle and bottom panels use *ϵ* = 10^*−*1^ and *ϵ* = 2, respectively. Black curves are trajectories; blue dashed lines are setpoints.

Figure 3 then relaxes the operating-regime assumptions in (12)–(13) one at a time. Unless otherwise stated, all parameters are identical to those in the middle panel of Fig. 2; in particular, *ϵ* = 0.1. In the top panel, 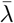 and 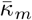 are reduced together so that *θ* remains constant, while *K*_*s*_ = *K*_*m*_ = 10 keeps the promoters near their linear regime; thus the main effect is the loss of sRNA-dominated mRNA removal. For the tunings associated with the idealized setpoints *r*_ss_ = 10, 1, 10^−1^, the quantity 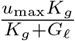 in (36) is approximately 0.517, 0.655, 1.302, respectively. Hence the tunings for *r*_ss_ = 10 and *r*_ss_ = 1 satisfy (36), whereas the tuning for *r*_ss_ = 10^−1^ does not. In the bottom panel, *K*_*s*_ and *K*_*m*_ are reduced to test loss of promoter linearity. In that case coexistence can still persist, but the steady-state ratio is no longer selected by *θ* alone, so the simple tuning rule (17) no longer predicts the actual coexistence ratio of the full implementation model.

**Fig. 3.**
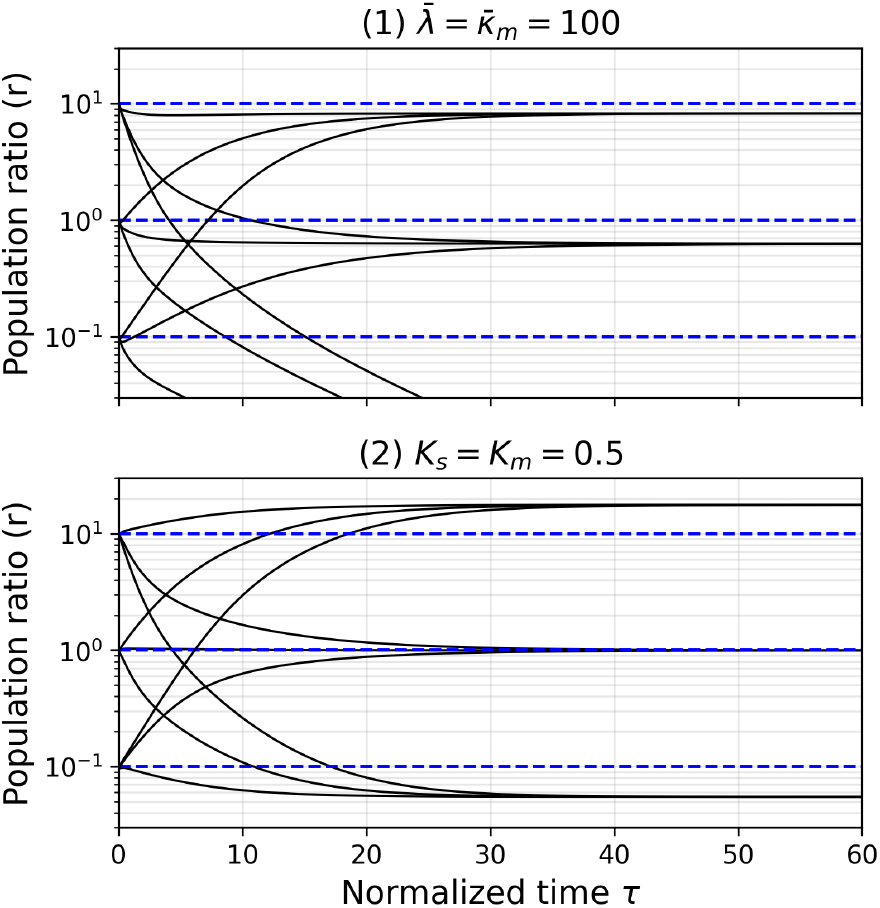
Violating either sRNA-dominance or promoter linearity breaks the idealized ratio-selection rule. Both panels simulate the full implementation model (18) at *ϵ* = 0.1, and should be compared with the middle panel of Fig. 2. The idealized design values are *θ* ∈ {0.05, 0.5, 5}, corresponding to the target setpoints *r*_ss_ ∈ {10^*−*1^, 10^0^, 10^1^}. In each panel, 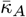 is chosen by matching the reduced full-system parameter *θ* to the corresponding idealized value; for these paired perturbations this gives 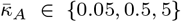 in Both panels. Common settings in both panels: 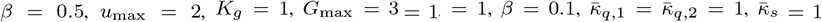, 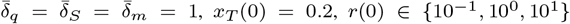, and initial states *x*_*s*_(0) = *x*_*T*_ (0)*r*(0)*/*(1 + *r*(0)), *x*_*f*_ (0) = *x*_*T*_ (0)*/*(1 + *r*(0)), *q*_*s*_(0) = *q*_*f*_ (0) = *S*(0) = *m*_*A*_(0) = *A*(0) = 0. Top panel: 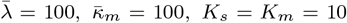 so sRNA-mediated degradation is weakened relative to the nominal design while the promoters remain near their linear regime. Bottom panel: 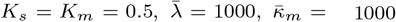, so the promoter maps are no longer well approximated by their linear regime. Black curves are trajectories; blue dashed lines are the idealized setpoints.

## VII. Conclusion

This paper developed a control-theoretic framework for programmable composition regulation in a two-strain consortium. The idealized ratio-feedback law (16) yields a tunable coexistence ratio through (17), and the full implementation model (18) realizes this behavior through quorum sensing, sRNA-mediated comparison, and ppGpp-driven growth actuation. Using singular perturbation theory, we showed that, for sufficiently small *ϵ* and under suitable operating conditions, the full implementation inherits the coexistence equilibrium and local stability of the reduced model.

Figure 2 shows that the full implementation reproduces the idealized closed-loop behavior when the reduction assumptions hold, whereas relaxing these assumptions degrades transient performance and can ultimately eliminate coexistence. Weakening sRNA-dominated mRNA removal causes loss of coexistence, whereas loss of promoter linearity mainly breaks the direct mapping between the design parameter and the steady-state ratio.

The main design conditions are that *ϵ* be small, the sensing promoters operate in the linear regime (12), sRNA-mediated mRNA removal dominate basal decay (13), and ppGpp-mediated actuation satisfy (36). The dominant controller timescale is likely set by RelA^+^ turnover, while the broader physiological effects of ppGpp remain an important practical consideration. Unlike the ratiometric design in [16], the controller is purely sRNA-based and avoids the additional ribosomal load associated with protein-mediated degradation, making it more closely aligned with the RNA-based design philosophy of [14].

Experimental validation could proceed in stages. First, the ratiometric sensor can be isolated by replacing RelA^+^ with a fluorescent reporter and manually varying consortium composition; the reporter should scale with the imposed ratio *x*_*s*_*/x*_*f*_, validating the sensing module. Second, the actuator timescale should be characterized separately. While previous work showed that RelA^+^ expression can tune growth rate [17], its turnover may dominate *ϵ*. Figure 2 suggests that even for *ϵ* = *O*(1), regulation is preserved, albeit with slower oscillatory transients. Degradation tags could potentially reduce this timescale, provided they do not compromise RelA^+^ function. Finally, the complete closed-loop circuit should be tested in continuous culture to verify that varying *θ* shifts the coexistence ratio according to (17). Future work should examine robustness more systematically, including sensitivity of steady-state and transient behavior to circuit and host parameters [19], [20]. Another natural extension is to relax the fast quorum-sensing assumption by including the quorum-sensing states in the slow subsystem. It would also be useful to study hybrid feedback-feedforward architectures, compare this controller with other growth-rate actuators, and extend the framework to larger consortia.

## VIII. AI Statement

ChatGPT (version 5.2) was used to assist in streamlining and editing portions of this manuscript and in developing code for the numerical simulations. The authors reviewed and validated all content.

## IX. Acknowledgments

This work was supported by internal research funding at the Johns Hopkins Applied Physics Laboratory.

